# Structural diversity and stress regulation of the plant immunity-associated CALMODULIN-BINDING PROTEIN 60 (CBP60) family of transcription factors in *Solanum lycopersicum* (tomato)

**DOI:** 10.1101/2022.12.06.519278

**Authors:** Vanessa Shivnauth, Sonya Pretheepkumar, Eric Marchetta, Keaun Amani, Christian Danve M. Castroverde

**Author notes:** These authors contributed equally (Sonya Pretheepkumar, Eric Marchetta).

## Abstract

Cellular signalling generates calcium (Ca^2+^) ions, which are ubiquitous secondary messengers decoded by calcium-dependent protein kinases, calcineurins, calreticulin, calmodulins (CAMs) and CAM-binding proteins. Previous studies in the model plant *Arabidopsis thaliana* have shown the critical roles of the CAM-BINDING PROTEIN 60 (CBP60) protein family in plant growth, stress responses and immunity. Certain CBP60 factors can regulate plant immune responses, like pattern-triggered immunity, effector-triggered immunity, and synthesis of major plant immune-activating metabolites salicylic acid (SA) and N-hydroxypipecolic acid (NHP). Although homologous CBP60 sequences have been identified in the plant kingdom, their function and regulation in most species remain unclear. In this paper, we specifically characterized 11 members of the CBP60 family in the agriculturally important crop tomato (*Solanum lycopersicum*). Protein sequence analyses revealed that three CBP60 homologs have the closest amino acid identity to *Arabidopsis* CBP60g and SARD1, master transcription factors involved in plant immunity. Strikingly, AlphaFold deep learning-assisted prediction of protein structures highlighted close structural similarity between these tomato and *Arabidopsis* CBP60 homologs. Conserved domain analyses revealed that they possess CAM-binding domains and DNA-binding domains, reflecting their potential involvement in linking Ca^2+^ signalling and transcriptional regulation in tomato plants. In terms of their gene expression profiles under biotic (*Pseudomonas syringae* pv. *tomato* DC3000 pathogen infection) and/or abiotic stress (warming temperatures), five tomato *CBP60* genes were pathogen-responsive and temperature-sensitive, reminiscent of *Arabidopsis CBP60g* and *SARD1*. Overall, we present a genome-wide identification of the CBP60 gene/protein family in tomato plants, and we provide evidence on their regulation and potential function as Ca^2+^-sensing transcriptional regulators.

## Introduction

Calcium is required for plant growth, development, and immunity (Hepler, 2005; Tian et al., 2020). Calcium ions in plant cells serve as intracellular messengers to elicit responses to different abiotic and biotic stressors (Knight, 2000; Köster et al., 2022; Xu et al., 2022). One of the earliest plant immune responses following pathogen recognition is a rapid influx of calcium ions into the cytosol (Moeder et al., 2019; Tian et al., 2019; Hilleary et al., 2020; Thor et al., 2020). Proteins such as calmodulin (CAM) bind calcium, and these calcium-binding proteins then alter their conformation and catalytic activity resulting in signal transduction (Yang and Poovaiah, 2003; DeFalco et al., 2009). CAM is a highly studied eukaryotic protein that interacts with numerous target proteins (Bouché et al., 2005; Kim et al., 2009). For example, CAM interacts with and activates certain CAM-binding transcription factors involved in immune responses, like CALMODULIN-BINDING TRANSCRIPTION ACTIVATOR 3 (CAMTA3; Du et al., 2009) and CALMODULIN-BINDING PROTEIN 60-LIKE G (CBP60g; Wang et al., 2009; Zhang et al., 2010; Sun et al., 2022).

CBP60g is a member of the CBP60 protein family (Reddy et al., 2002; Wang et al., 2009; Truman et al., 2013; Amani et al., 2022; Zheng et al., 2022) and serves as a key transcriptional regulator for SA biosynthetic genes *ISOCHORISMATE SYNTHASE 1 (ICS1*) and *AVRPPHB SUSCEPTIBLE 3 (PBS3;* Zhang et al., 2010; Wang et al., 2009; Sun et al., 2015; Kim et al., 2022). Like CBP60g, another CBP60 protein family member SYSTEMIC ACQUIRED RESISTANCE 1 (SARD1) plays a partially redundant role in SA biosynthesis (Zhang et al., 2010; Wang et al., 2011). Although SARD1 does not bind CAM (unlike CBP60g), it has been shown to be regulated by calcium sensor proteins like CALCIUM-DEPENDENT PROTEIN KINASE 5 (CPK5; Guerra et al., 2020). Apart from SA production, CBP60g and SARD1 also positively regulate systemic acquired resistance by controlling genes like *AGD2-LIKE DEFENSE RESPONSE PROTEIN 1 (ALD1), SYSTEMIC ACQUIRED RESISTANCE 1 (SARD4*) and *FLAVIN-DEPENDENT MONOOXYGENASE 1 (FMO1;* Sun et al., 2018; Shields et al. 2022), which are required for biosynthesis of the systemic immunity-activating metabolite *N*-hydroxypipecolic acid (NHP; Chen et al., 2018; Hartmann et al., 2018; Huang et al., 2020; Zeier, 2021).

CBP60g and SARD1 are two of eight homologous proteins of the CBP60 family in *Arabidopsis* and are strongly inducible by pathogen infection (Wang et al., 2009; Zhang et al., 2010; Wang et al., 2011; Truman et al., 2013). In *A. thaliana* plants, other CBP60 family members include CBP60a, which is a CAM-binding negative regulator of immunity as CBP60a mutations reduced pathogen growth (Truman et al., 2013; Lu et al., 2018). As another member of the *Arabidopsis* CBP60 family, CBP60b functions as a positive regulator for both cell surface and intracellular immune receptors (Huang et al., 2021; Li et al., 2021). CBP60b has also been found to bind the *SARD1* promoter region, which suggests that it could regulate *SARD1* expression (Huang et al., 2021). CBP60c and CBP60d mutations have small significant effects on plant disease resistance, while the effects of CBP60d and CBP60e on plant immunity seem negligible (Truman et al., 2013).

Importantly, the temperature-sensitivity of the SA biosynthetic pathway (Huot et al., 2017; Castroverde and Dina, 2021) is due to the temperature-downregulation of *CBP60g* and *SARD1* (Kim et al., 2022). *CBP60g* and *SARD1* gene expression can be induced by pathogens or pathogen-associated molecular patterns (Wang et al., 2009; Zhang et al., 2010; Wang et al., 2011), but induced expression is suppressed when temperatures increase (Kim et al., 2022). Remarkably, constitutive expression of *CBP60g* or *SARD1* can restore not only SA biosynthesis at warm temperature but also other drivers of the plant immune system (Kim et al., 2022). *CBP60g* and *SARD1* gene expression are tightly regulated, with transcription factors TGA1 and TGA4 acting as positive regulators (Sun et al., 2018) and CAMTA proteins as negative regulators (Sun et al., 2020).

Because of the central importance of CBP60g and SARD1 in plant immune resilience to a warming climate, it is imperative that functional orthologs are investigated in other plant species, especially agriculturally important crops. Although a recent study reported orthologs of CBP60g and SARD1 in tobacco plants (Takagi et al., 2022), the function and regulation of CBP60 proteins in other plant species have yet to be investigated. We recently identified CBP60 homologs across various representative taxa in the plant kingdom (Amani et al., 2022); however, whether gene expression trends observed in *Arabidopsis* are conserved in other plants remain unclear. In this study, we report the identification of 11 homologous *CBP60 (SlCBP60*) genes in tomato plants (*Solanum lycopersicum*). Our analyses show that SlCBP60-1, 8 and 11 are the closest sequence and structural homologs to *Arabidopsis* CBP60g and SARD1. In addition, we show that biotic stress (pathogen infection) and abiotic stress (elevated temperature) differentially regulate the 11 *SlCBP60* genes, with observed variation in pathogen-responsiveness and temperature-vulnerability.

## Materials and Methods

### Protein sequence analyses

Protein IDs of the 11 tomato (*S. lycopersicum*) CBP60 homologs or SlCBP60 were obtained from Gramene (https://www.gramene.org/; Tello-Ruiz et al., 2021). Amino acid sequences were then exported from the Sol Genomics Network (https://solgenomics.net/; Fernandez-Pozo et al., 2015). SlCBP60 protein sequences were analyzed for amino acid similarity/clustering using Molecular Evolutionary Genetics Analysis (MEGA) Bioinformatics (Kumar et al., 1994), where they were built into a protein sequence alignment using the MUSCLE algorithm (Edgar, 2004). A dendrogram of the 11 SlCBP60g homologs was constructed as a Neighbor-Joining Tree together with the reference *A. thaliana* SARD1 and CBP60g protein sequences obtained from The *Arabidopsis* Informatics Resource/TAIR (https://www.Arabidopsis.org/; Lamesch et al., 2012). In addition, SlCBP60 protein sequences were analyzed for putative CAM-binding domains through Pfam (http://pfam.xfam.org/null; Mistry et al., 2021) and putative DNA-binding domains through DP-Bind (http://lcg.rit.albany.edu/dp-bind/; Hwang et al., 2007). Finally, candidate SlCBP60 phosphosites were determined by comparing with confirmed AtCBP60g and AtSARD1 phosphosites compiled in the qPTMPlants website (http://qptmplants.omicsbio.info/; Xue et al., 2022).

### AlphaFold protein structural prediction and hierarchical clustering

Protein structures of the 11 tomato SlCBP60 homologs were predicted using the ColabFold: AlphaFold2 with MMseqs2 model (https://github.com/sokrypton/ColabFold; Jumper et al., 2021; Mirdita et al., 2022). Structures were predicted by inputting their corresponding amino acid sequences to the model using the default configuration. After the protein structures were predicted through AlphaFold2, the model outputted 5 structures ranked based on the model’s confidence in each structure. For each tomato SlCBP60 protein, we examined the highest-ranked structure automatically computed by AlphaFold2. To visualize the protein structures, the resulting PDB file formats were uploaded to the RCSB PDB website (https://www.rcsb.org/3d-view; Burley et al., 2019).

TM-score analyses to determine similarities between predicted protein structures were conducted through the Zhang Lab website (https://zhanggroup.org/TM-score/; Zhang and Skolnick, 2004). All 11 protein structures were inputted in PDB format, and TM-scores were compared with the other tomato SlCBP60 proteins and with the reference *Arabidopsis* proteins AtCBP60g and AtSARD1. The TM-scores were analyzed by hierarchical clustering using the NG-CHM Builder tool (https://build.ngchm.net/NGCHM-web-builder/; Ryan et al., 2019). Row and column ordering were set to “hierarchical clustering.” The distance metric used was “Euclidean” and the agglomeration setting was “average linkage.”

### Promoter analyses and transcription factor binding predictions

Upstream DNA sequences of the 11 *SlCBP60* genes were obtained using PlantPAN 3.0 (http://plantpan.itps.ncku.edu.tw/; Chow et al., 2019). The upstream and downstream coordinates of promoter transcription start site/5’UTR-End were set to X: 1000 and Y:100, for upstream and downstream of the gene, respectively. *SlCBP60* gene promoter sequences were then analyzed for nucleotide sequence similarity/clustering using Molecular Evolutionary Genetics Analysis (MEGA) Bioinformatics (Kumar et al., 1994). Putative transcription factors that bind to the 11 *SlCBP60* promoters were predicted using PlantPAN 3.0 using the Multiple Promoter Analysis tool. Unique and overlapping transcription factors were sorted using UpSetR to visualize interactions in a matrix layout (Conway et al., 2017).

### Plant materials and growth conditions

Tomato cultivar Castlemart seeds were kindly provided by Dr. Gregg Howe from Michigan State University (Li et al., 2004). Seeds were sterilized in 10% bleach solution for 15 minutes and washed five times with autoclaved water. Seeds were then hydrated with autoclaved water at room temperature (21°-23°C) overnight to facilitate imbibition. Afterwards, seeds were allowed to germinate on sterile 9-cm filter paper for 5 days under dark conditions. Germinated seeds were planted in pots (9.7cm x9.7cm) containing autoclaved soil (3 parts Promix PGX and 1 part Turface). Individual plants were initially fertilized with 100mL of MiracleGro solution (made with a ratio of 4 g of MiracleGro per 1 L of water). Tomato plants were grown at 23°C with a 12 hr light (100 ±20 umol m^-2^ s^-1^) and 12 hr dark cycle and 60% relative humidity. Plants were watered regularly and fertilized weekly with nutrient water (Hoagland and Arnon, 1950).

### Pathogen infection

For pathogen-induced gene expression analyses, one leaf from 4-week-old plants was infiltrated using a needleless syringe with either mock (0.25mM MgCl_2_) or *Pseudomonas syringae* pv. *tomato/Pst* DC3000 (OD600=0.001) as previously described in detail (Huot et al., 2017). Inoculated plants were incubated at either normal (23°C day/23°C night) or elevated temperature (32°C day/32°C) with 60% relative humidity and 100 ±20 umol m^-2^ s^-1^ light intensity. For systemic gene expression analyses, mature and healthy bottom leaflets of 3- to 4-week-old tomato plants were infiltrated with either mock (0.25mM MgCl_2_) or *Pst* DC3000 (OD600=0.02) based on a protocol by Holmes et al. (2019). Inoculated plants were incubated at normal temperature (23°C day/23°C night) with 60% relative humidity and 100 ±20 umol m^-2^ s^-1^ light intensity. Four individual plants were used as independent biological replicates per treatment.

### Gene expression analyses

Locally infected leaves were harvested at 24 hours after mock or pathogen treatment, while uninfected (upper) systemic leaflets were harvested at 48 hours after local treatment of lower leaflets. Gene expression levels were quantified based on a previously published protocol (Huot et al., 2017; Kim et al., 2022) with slight modifications. After tissue homogenization using the TissueLyser II (Qiagen), total RNA was extracted using the RNeasy Plant Mini Kit (Qiagen). Total RNA concentration and quality were measured using a Nanodrop (Thermo Fisher) or DeNovix Nanospec. The cDNA was synthesized using qScript cDNA super mix (Quantabio) based on manufacturers’ recommendations. Real-time quantitative polymerase chain reaction (qPCR) was performed using PowerTrack SYBR Green master mix (Life Technologies). Equivalently diluted mRNA without the qScript cDNA mix were used as negative controls. The resulting qPCR mixes were run using the Applied Biosystems QuantStudio3 platform (Life Technologies), and individual Ct values were determined for target genes and the internal control gene (*SlACT2*) (Dekkers et al., 2012). Gene expression values were reported as 2^-ΔCt^, where ΔCt is Ct_target gene_–Ct_*SlACT2*_. qPCR was carried out with three technical replicates for each biological sample. Primers used for qPCR are shown in Supplementary Table 1.

## Results

### Protein sequence analyses and phylogeny of the 11 SlCBP60 homologous proteins in tomato plants

Unlike CBP60 proteins in the model species *A. thaliana* (Reddy et al, 2002; Wang et al., 2009; Zhang et al., 2010; Wang et al., 2011; Wan et al., 2012; Truman et al., 2013), CBP60 proteins in *S. lycopersicum* (tomato) plants have remained uncharacterized. Using Gramene, we successfully identified 11 *CBP60* homologous genes in tomato: *Solyc01g100240* (*SlCBP60_1*), *Solyc02g079040* (*SlCBP60_2*), *Solyc03g113920* (*SlCBP60_3*), *Solyc03g113940* (*SlCBP60_4*), *Solyc03g113950* (*SlCBP60_5*), *Solyc03g113960* (*SlCBP60_6*), *Solyc03g113970* (*SlCBP60_7*), *Solyc03g119250* (*SlCBP60_8*), *Solyc07g006830* (*SlCBP60_9*), *Solyc10g009210* (*SlCBP60_10*) and *Solyc12g036390* (*SlCBP60_11*). There is one homologous *CBP60* gene each for chromosomes 1, 2, 7, 10 and 12, while there are six *CBP60* homologs in chromosome 3 alone. The SlCBP60 protein sequences are listed in Supplementary Table 2.

To shed light on potential function and diversification of the 11 tomato SlCBP60 proteins, we analyzed their primary amino acid sequence similarities. As shown in Figure 1A, phylogenetic analyses revealed two main clades of tomato SlCBP60 proteins. The first major clade had four subclades: (a) SlCBP60_8 and 11; (b) SlCBP60_2, 9 and 10; (c) SlCBP60g_1; and (d) SlCBP60g_3. The second major clade had two subclades: (a) SlCBP60g_4 and 5; and (b) SlCBP60g_6 and 7. We had built the reference CBP60 proteins (AtCBP60g and AtSARD1) from the model species *A. thaliana* into this protein phylogenetic analyses. Based on their amino acid identities, SlCBP60_1 is the closest homolog to *Arabidopsis* CBP60g, while *Arabidopsis* SARD1 is most directly related to SlCBP60_8 and SlCBP60_11.

**Fig. 1.**
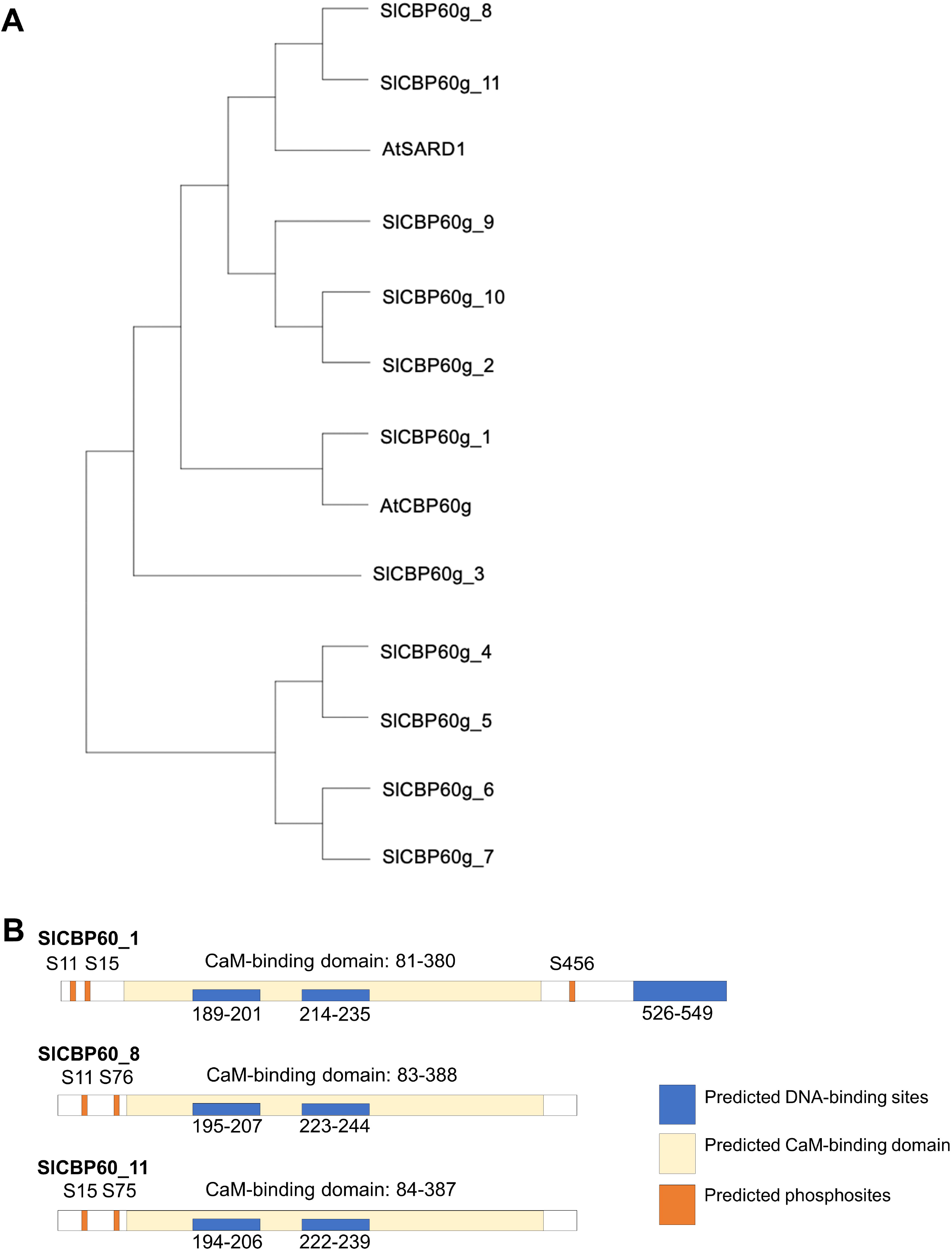
Sequence analyses of the tomato SlCBP60 proteins. (A) Tomato SlCBP60 sequences were obtained from Sol Genomics Network (https://solgenomics.net). *A. thaliana* sequences were obtained from TAIR (https://www.Arabidopsis.org/). Sequences were built into a protein sequence alignment using the MUSCLE algorithm on MEGA and a neighbour-joining tree was constructed. (B) Close tomato homologs to *Arabidopsis* CBP60g (SlCBP60_1) and SARD1 (SlCBP60_8 and SlCBP60_11) were further analyzed for putative CAM-binding domains (using Pfam) and DNA-binding sites (using DP-Bind). Conserved phosphosites are also indicated based on experimentally identified AtCBP60g and AtSARD1 phosphoserines (using qPTMPlants).

Having identified SlCBP60_1, 8 and 11 as the closest sequence homologs of AtCBP60g and AtSARD1, we performed functional domain analyses to confirm whether they possess the distinguishing hallmarks of CBP60 family transcription factors. As shown in Figure 1B, all three SlCBP60 paralogs have predicted CAM-binding domains, suggesting their mechanistic link to plant calcium signalling. Putative DNA-binding residues were also detected in the three proteins, with two proximal DNA-binding domains within the CAM-binding domain. Additionally, the longer SlCBP60_1 protein also contained a third DNA-binding domain in its C-terminus. This is consistent with AtCBP60g being longer than its AtSARD1 paralog (Zhang et al., 2010; Wang et al., 2011). Finally, by examining protein phosphosites on qPTMPlants, we determined conserved phosphoserine residues in the putative tomato orthologs. In SlCBP60_1, the Ser11, Ser15 and Ser456 residues correspond to experimentally determined phosphosites in AtCBP60g (Ser8, Ser11 and Ser450; Xue et al., 2022). On the other hand, SlCBP60_8 Ser11/76 and SlCBP60_11 Ser15/75 residues were consistent with the AtSARD1 phosphosites (Ser12 and Ser77; Xue et al., 2022).

### Structural similarity analyses of the tomato SlCBP60 proteins

Three-dimensional protein structures are important to understand protein function. To predict structures of the 11 tomato SlCBP60 proteins, corresponding amino acid sequences were used as inputs to ColabFold, which uses AlphaFold2 with MMseqs1 model (Jumper et al., 2021; Mirdita et al., 2022). The AlphaFold model outputted and ranked five structures based on the model’s confidence in each structure. The highest-ranked predicted protein structures are shown in Figure 2A and Supplementary Data 1.

**Fig. 2.**
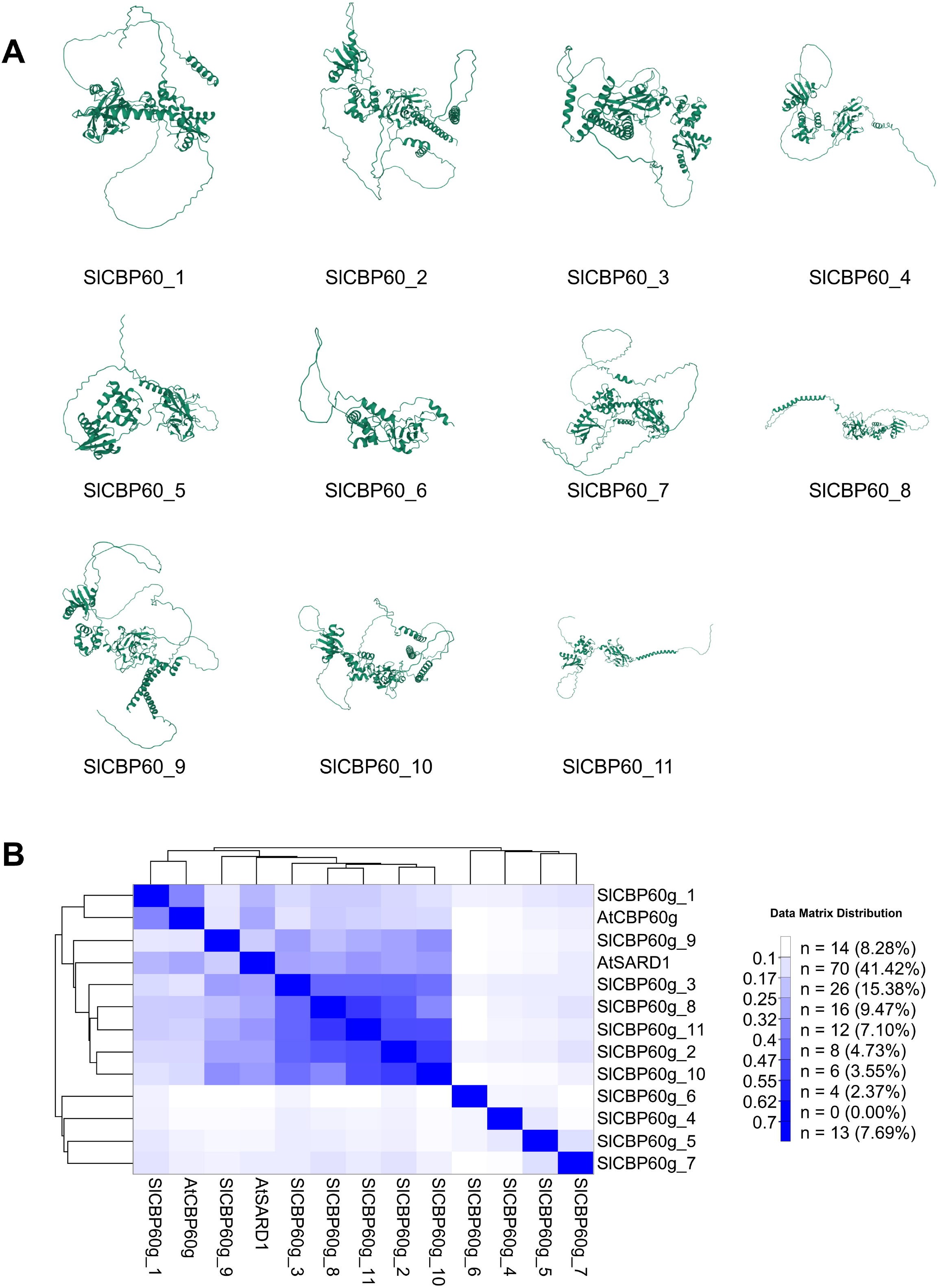
Structural similarity of AlphaFold deep learning-predicted SlCBP60 protein structures in tomato. A) Protein structures of the 11 tomato SlCBP60 homologs were predicted using AlphaFold2 with MMseqs2 model through the ColabFold Notebook (https://github.com/sokrypton/ColabFold). The best-ranked structure for each protein was visualized using the RCSB Protein Data Bank (https://www.rcsb.org/3d-view). (B) Hierarchical clustering of 11 tomato SlCBP60 protein structures was performed. Pairwise TM-scores were determined for all SlCBP60 proteins and the reference *Arabidopsis* proteins AtCBP60g and AtSARD1 on the Zhang Lab website (https://zhanggroup.org/TM-score/; Zhang et al., 2004). The TM-scores were then analyzed by hierarchical clustering using the NG-CHM Builder tool (https://build.ngchm.net/NGCHM-web-builder/).

To quantitatively determine structural similarity among the proteins, TM-scores were obtained to assess topological similarity of protein structures. Pairwise TM-score analyses were performed between each SlCBP60 protein and the reference *Arabidopsis* AtCBP60g and AtSARD1 proteins (Figure 2B; Supplementary Table 3). TM-scores with a value of 1.0 indicate perfect identity between two structures, while scores below 0.17 indicate unrelated proteins (Zhang et al., 2004). Based on the TM-score values and the structural similarity hierarchical clustering, SlCBP60_1 bears the most similar protein folding as AtCBP60g (consistent with the sequence analyses in the previous section). Also in agreement with the Figure 1A dendrogram, SlCBP60_8 and 11 structurally cluster together with AtSARD1. It is important to note that three other tomato proteins share structural similarity with AtSARD1 in this cluster (SlCBP60_2, 3 and 10). Finally, the distinct clade of distantly related SlCBP604, 5, 6 and 7 sequences (Figure 1A) also formed their own structural cluster in Figure 2B.

### Gene expression analyses of the tomato *SlCBP60* genes under bacterial pathogen infection at elevated temperature

Protein function can be potentially inferred based on their expression profiles. In *Arabidopsis, AtCBP60g* and *AtSARD1* gene expression in terms of transcript levels are induced by pathogens like *Pst* DC3000 at normal ambient temperatures, consistent with their central regulatory roles in the plant immune system (Wang et al., 2009; Zhang et al., 2010; Wang et al., 2011; Sun et al., 2015). These two master immune transcription factors are also critical for the vulnerability of plant immune responses under warm temperatures, since *AtCBP60g* and *AtSARD1* transcript levels are suppressed at elevated temperature (Kim et al., 2022).

To determine how both biotic (pathogen infection) and abiotic stresses (warm temperature) regulate tomato *SlCBP60* gene expression, total RNA samples were collected from tomato leaves after mock and pathogen treatments under both normal and elevated temperatures. As shown in Figure 3, RT-qPCR analyses indicated that *SlCBP60-2, 6, 8, 9* and *11* genes were induced after pathogen infection, while *SlCBP60-1, 3, 4, 5, 7* and *10* exhibited pathogen-unresponsive gene expression. It is important to note that we sometimes observed pathogen-induced *SlCBP60_1* gene expression in some but not all samples. In terms of temperature-sensitivity, all pathogen-induced genes exhibited temperature-sensitivity, while those not regulated by pathogen infection were resilient to temperature changes. Remarkably, the phylogenetically distant clade of *SlCBP60-4, 5, 6* and *7* generally had the lowest levels of gene expression.

**Fig. 3.**
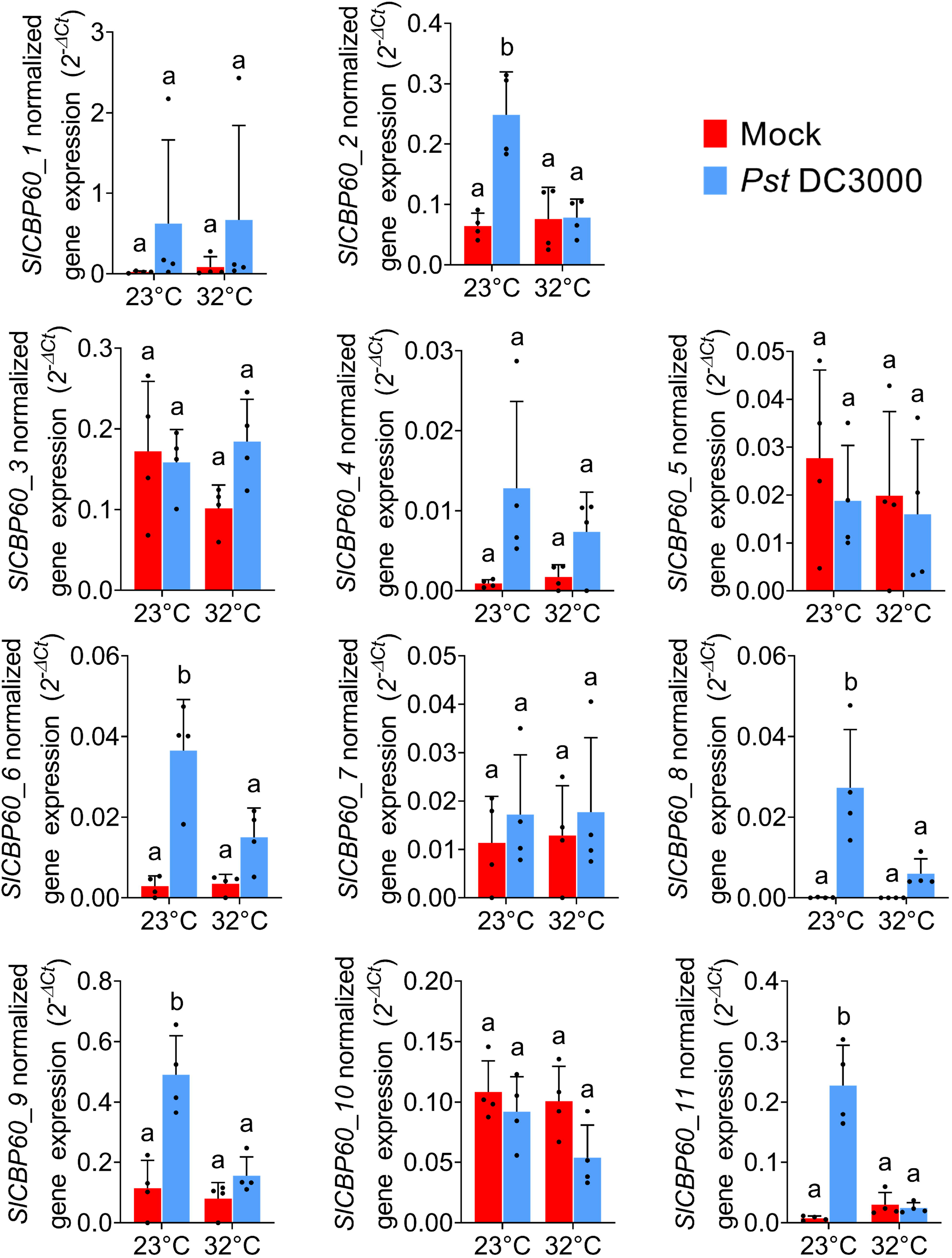
Gene expression analyses of tomato *CBP60* genes after pathogen infection under normal and elevated temperatures. Leaves of three- to four-week-old tomato plants were collected 1 day after syringe-infiltration with mock solution (0.25 mM MgCl_2_) or *Pst* DC3000 (OD600=0.001). Total RNA samples were extracted and used as templates for RT-qPCR with primers specific for *SlCBP60_1* to *SlCBP60_11*. Results show the mean gene expression value (relative to *SlActin2*) ± standard deviation of four biological replicates (n=4) of one representative experiment. Statistical significance was determined using a one-way ANOVA with Tukey’s honestly significant difference test (*p* < 0.05). Treatments with statistically significant differences are indicated by different letters. The experiment was performed three to four times with reproducible results.

### Systemic expression of the tomato *CBP60* genes after immune elicitation

In *Arabidopsis* plants, *AtCBP60g* and *AtSARD1* are induced systemically during systemic acquired resistance (Zhang et al., 2010). To elucidate how local immune elicitation also regulates systemic tomato *SlCBP60* gene expression, gene expression profiles of the 11 *SlCBP60* homologs were measured systemically after local infection with *Pst* DC3000. Relative transcript levels were compared between mock-treated and SAR-activated tomato plants as shown in Figure 4. It is evident that none of the *SlCBP60* genes exhibited statistically significant systemic induction after pathogen infection. These included genes that were induced locally after pathogen infection – *SlCBP60_2, 6, 8, 9* and *11*. Consistent with the results in the previous section, basal expression levels were highest for the constitutively expressed *SlCBP60_3* and *9* genes and were lowest for *SlCBP60_4, 5, 6, 7, 8* and *11*.

**Fig. 4.**
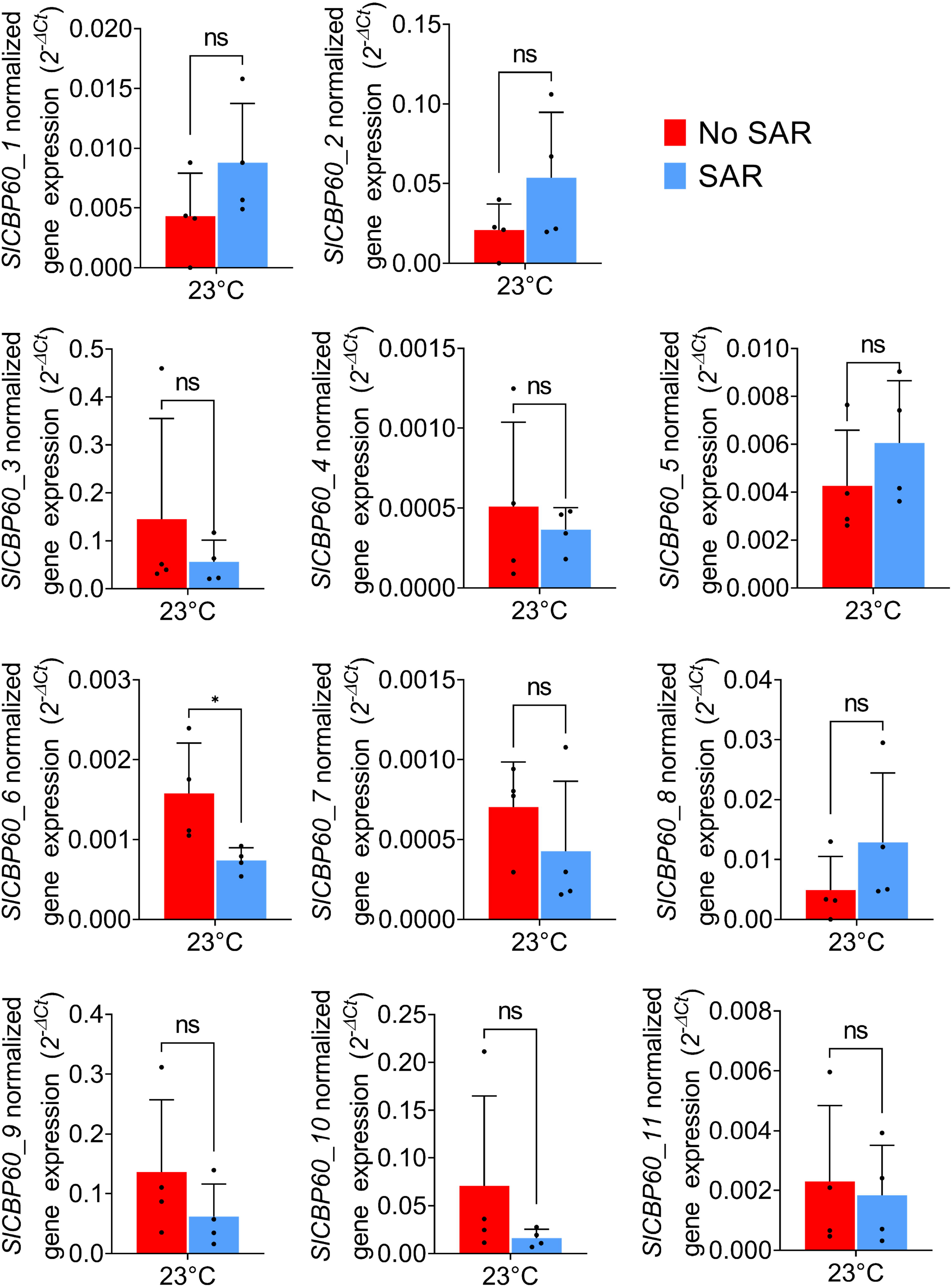
Gene expression analyses of tomato *SlCBP60* genes after systemic immune elicitation. Upper systemic leaflets of three- to four-week-old tomato plants were collected 2 days after infiltrating lower leaflets with mock solution (0.25 mM MgCl_2_) or *Pst* DC3000 (OD600=0.02). Total RNA samples were extracted and used as templates for RT-qPCR with primers specific for *SlCBP60_1* to *SlCBP60_11*. Results show the mean gene expression value (relative to *SlActin2*) ± standard deviation of four biological replicates (n=4) of one representative experiment. Statistical significance was determined using a pairwise t-test (*p* < 0.05), with asterisks (*) indicating statistically significant differences and “ns” indicating non-significant differences. The experiment was performed two times with reproducible results.

### In silico analyses of the tomato *SlCBP60* promoter regions

To characterize overall similarity and clustering in the tomato *SlCBP60* gene promoter sequences, we performed similarity clustering of their upstream DNA sequences using MEGA. As shown in Figure 5A, the phylogenetic tree for the 11 tomato *SlCBP60* gene promoter sequences resulted in two major clades. The first clade had four subclades: (a) *SlCBP60_1* and *6* promoters; (b) *SlCBP60_2* and *3* promoters; (c) *SlCBP60_8* and *10* promoters; and (d) *SlCBP60_7* and *11* promoters. The second clade had 3 members: *SlCBP60_4, 5* and *9* promoters. It is important to note that each clade/subclade consisted of both temperature-sensitive pathogen-induced genes and temperature-resilient constitutively expressed genes.

**Fig. 5.**
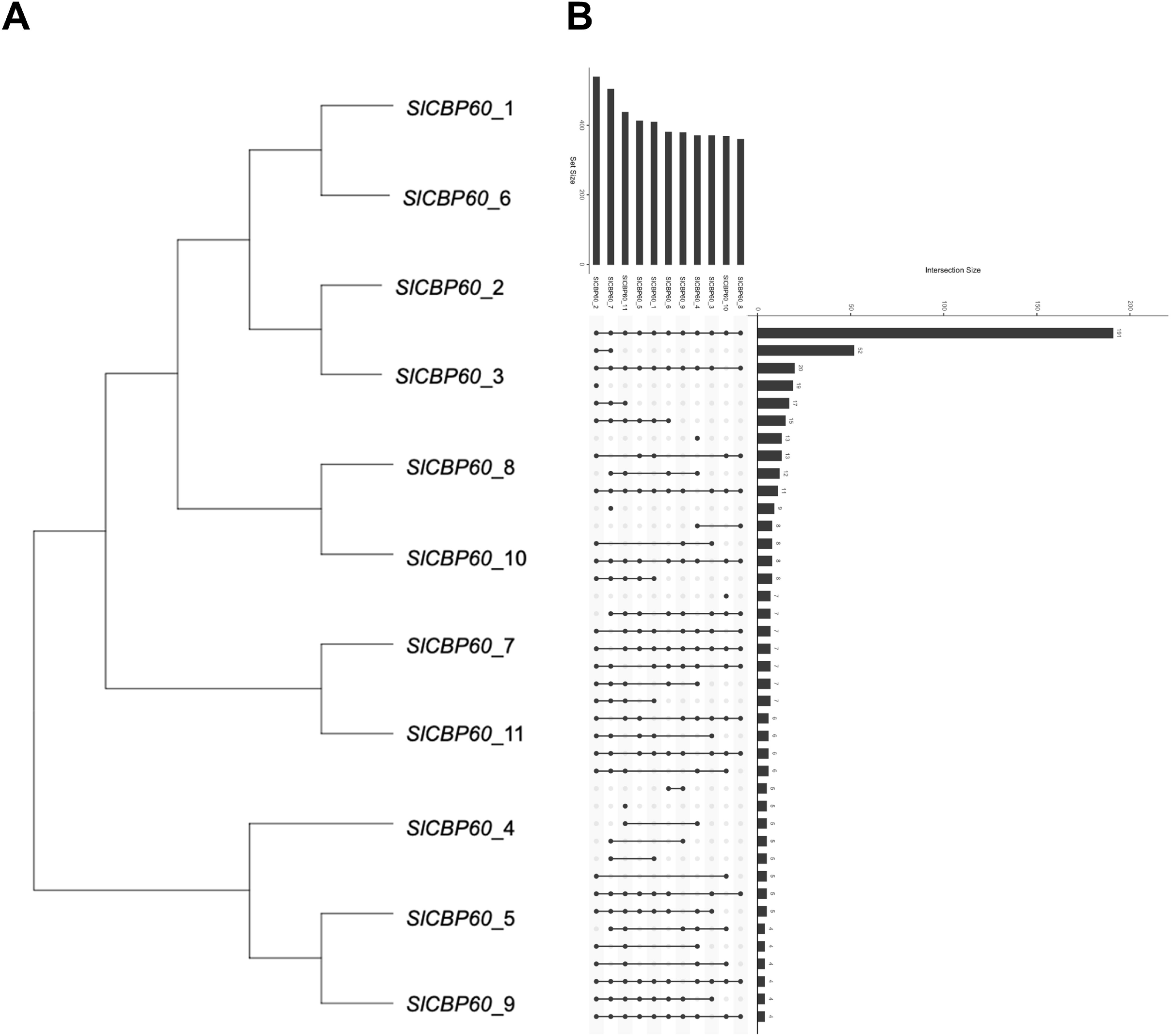
Sequence analyses and transcription factor binding predictions of the 11 tomato SlCBP60 gene promoter sequences. (A) *SlCBP60* promoter sequences were downloaded from the PlantPAN 3.0 website (http://plantpan.itps.ncku.edu.tw/) and then analyzed for similarity/clustering using MEGA. (B) Putative transcription factors that bind to the 11 *SlCBP60* upstream sequences were determined using PlantPAN 3.0. Unique and overlapping transcription factors were sorted using UpSetR to visualize set interactions in a matrix layout (Conway et al., 2017). The sets are ordered by intersection size, which indicates the number of transcription factors shared between the tomato *SlCBP60* gene promoter sequences. Sets with an exclusive intersection are filled with a dark circle and sets with no exclusive intersection are indicated by a light-gray circle.

Subsequently, a Multiple Promoter Analysis was performed in PlantPAN 3.0 to predict putative transcription factors that could bind the 11 *SlCBP60* promoter regions (Supplementary Table 4). The predicted transcription factors were visualized with UpsetR as shown in Figure 5B. From this analysis, *SlCBP60_1* to *11* shared 191 common transcription factors. The second intersection size was shared between the pathogen-induced *SlCBP60_2* gene and constitutively expressed *SlCBP60_7* gene (52 common transcription factors). Next, all genes except the constitutively expressed *SlCBP60_10* gene shared another 20 common transcription factors. The pathogen-induced *SlCBP60_2* gene had 19 unique transcription factors, while it shared another 17 transcription factors uniquely with *SlCBP60_7* and *11*. Additionally, *SlCBP60_1, 2, 5, 6, 7* and *11* shared 15 unique transcription factors. Independently, the constitutively expressed *SlCBP60_4* and *SlCBP60_7* genes had 13 and 9 unique transcription factors, respectively. There were 13 common transcription factors for *SlCBP60_1, 2, 5, 8* and *10*, while *SlCBP60_4, 6, 7* and *11* shared 12 common transcription factors. All 11 *SlCBP60* genes except *SlCBP60_4* commonly shared 11 common transcription factors. Finally, other *SlCBP60* promoter interaction sets had less than 10 overlapping transcription factors.

## Discussion

In this paper, we have successfully identified and characterized 11 CBP60 family members in tomato plants. Unlike CBP60 proteins in the model species *Arabidopsis thaliana* (Reddy et al., 2002; Wang et al., 2009; Truman et al., 2013; Amani et al., 2022; Zheng et al., 2022), CBP60 proteins in tomato and other species have remained unexplored. First, phylogenetic and structural analyses were conducted for the 11 SlCBP60 proteins (Figures 1–2). Second, expression profiles of the 11 *SlCBP60* genes were determined after local and systemic immune elicitation with the model bacterial pathogen *Pst* DC3000 under different temperatures (Figures 3–4). Third, putative transcription factors that bind the *SlCBP60* gene promoters were predicted to potentially explain the differential regulation of these genes under biotic and abiotic stress (Figure 5).

Phylogenetic analyses revealed two major clades of the 11 tomato SlCBP60 proteins. The first clade clustered with the reference *Arabidopsis* AtCBP60g and AtSARD1 proteins. In particular, SlCBP60_1 has the highest amino acid identity to AtCBP60g, while SlCBP60_8 and 11 are closest phylogenetically to AtSARD1. High amino acid sequence conservation was observed in the middle region of the 11 SlCBP60 proteins, with most sequence differences observed in their C-terminal regions. Our sequence-guided ortholog analyses were further validated by AlphaFold-predicted protein structures and TM-score analyses for topological similarity (Zhang and Skolnick, 2004; Jumper et al., 2021; Mirdita et al., 2022). Based on TM-scores, SlCBP60_1 and AtCBP60g exhibit close structural similarity, while SlCBP60_8, SlCBP60_11 and AtSARD1 belong to another cluster of structurally similar proteins. Interestingly, other SlCBP60 proteins (2, 3, 9 and 10) also share close structural similarity to AtSARD1. The fact that the MEGA-generated phylogenetic tree (Figure 1) and TM-score-based structural clustering (Figure 2) did not perfectly mirror each other suggests that similarities not evident from primary amino acid sequences alone can be revealed by tertiary structural analyses. What is evident is that the distinct sequence subclade of SlCBP60_4, 5, 6 and 7 also forms a distinct and distantly related structural cluster.

Previous research in the highly studied model species *A. thaliana* demonstrated that the CBP60 family has a highly conserved domain in the central region (Zhang et al., 2010), which is congruent with our Pfam-predicted CAM-binding domains in all three close SlCBP60 homologs (Figure 1). AtCBP60g protein also has a confirmed CAM-binding domain located near the N-terminus (Wang et al., 2009), but we were not able to determine this in silico for SlCBP60_1. Remarkably, CAM-binding domains were predicted in SlCBP60_8 and 11 even though their closest homolog AtSARD1 cannot bind CAM (Zhang et al., 2010; Wang et al., 2011). It is important to note that a SARD1 ortholog in *Nicotiana tabacum* (NtSARD1) can bind CAM, indicating differential post-translational regulation of these proteins depending on the species. Furthermore, a previous study has shown a transcription activation domain in the AtCBP60g protein at residues 211-400 (Qin et al., 2018). These nicely fit within the predicted DNA-binding domains in SlCBP60_1 (residues 214-235), SlCBP60_8 (residues 223-244) and SlCBP60_11 (residues 222-239). Altogether, SlCBP60_1,8 and 11 may be the functional tomato orthologs of the *Arabidopsis* CBP60g and SARD1 proteins, which are master transcription factors controlling SA biosynthesis and immunity. However, further genetic confirmation is needed. Finally, we found conserved serine residues in these three proteins that correspond with the experimentally determined phosphosites in AtCBP60g and AtSARD1 based on previous studies (Xue et al., 2022; Sun et al., 2022). It would be interesting to explore whether these putative SlCBP60 phosphosites are also phosphorylated after immune elicitation and then to identify kinases and/or phosphatases responsible for this dynamic phosphorylation.

After our sequence- and structure-guided analyses of the tomato SlCBP60 proteins, we set out to determine how *SlCBP60* gene expression is regulated by stress conditions. In particular, we were curious to characterize which tomato genes would exhibit the same pathogen-induced expression of the *Arabidopsis AtCBP60g* and *AtSARD1* genes, which are vulnerable to suppression at elevated temperatures (Kim et al., 2022). Based on RT-qPCR analyses of these genes after *Pst* DC3000 pathogen infection at 23°C and 32°C (Figure 3), we discovered that *SlCBP60-2, 6, 8, 9* and *11* show temperature-modulated pathogen-induced gene expression that reflect transcriptional trends in *AtCBP60g* and *AtSARD1*. Interestingly, the closest sequence and structural homolog of AtCBP60g in tomato (SlCBP60_1) showed temperature-resilient constitutive levels of gene expression. Constitutively expressed genes could be further classified into those with low (*SlCBP60-1, 4, 5* and *7*) or high basal levels (*SlCBP60_3* and *10*), potentially reflecting differential functional, spatial and/or temporal regulation of these genes. In addition to local pathogen induction, *Arabidopsis AtCBP60g* and *SARD1* can be induced in uninfected distal tissues during systemic acquired resistance (Zhang et al., 2010; Shields et al., 2022). However, we did not observe systemically induced expression of any of the 11 *SlCBP60* genes after local immune elicitation with the virulent bacterial pathogen *Pst* DC3000 (Figure 4). This could suggest differential regulation of *CBP60* genes by mobile systemic immune signals between tomato and *Arabidopsis* plants.

Finally, to mechanistically link gene expression profiles with upstream transcriptional regulators, we analyzed promoter sequences of the 11 tomato *SlCBP60* genes and then predicted their putative transcription factors. Our findings demonstrate partial correlation between promoter sequence similarity and predicted transcription factor sets. For example, *SlCBP60_2* and *7* share 52 unique common transcription factors, and their promoter sequences cluster phylogenetically in a major clade. However, there were some unexpected results, such as *SlCBP60_1,2,5,8* and *10* sharing 13 transcription factors, even though their promoter sequences are distributed all over separate clades or subclades. Surprisingly, little correlation is observed between immunity-elicited gene expression profiles and shared transcription factors. *SlCBP60_2, 6, 8, 9* and 11 are pathogen-induced genes, but their promoter sequences are distributed across five distinct subclades, and we did not identify transcription factors that are shared exclusively among them. We also did not identify any transcription factor that are only shared among the constitutively expressed genes (*SlCBP60_1, 3, 4, 5, 7* and *10*). In the future, it may be necessary to investigate beyond the distal (short-distance) promoter regions. There may be non-local (distal) enhancer regions (Andersson and Sandelin, 2020) and/or three-dimensional chromatin architecture (Jerkovic and Cavalli, 2021) that could account for the differential regulation of the tomato *SlCBP60* gene family. In general, regulatory transcription factors not only rely on short-distance/proximal promoter regions, but they can be influenced by long-distance enhancer regions as well (Dong et al., 2017; Li et al., 2019; Yan et al., 2019).

Overall, our research has highlighted the structural and regulatory diversity of the 11 *SlCBP60* genes and their encoded proteins in tomato plants. We have identified candidate orthologs for further functional characterization. Our genome-wide structural and gene expression analyses have started to shed light on the potential involvement of these tomato SlCBP60 proteins in linking calcium signalling and transcriptional regulation of plant immunity.

## Funding and Acknowledgements

We would like to thank: Dr. Gabriel Moreno-Hagelsieb for helping with the hierarchical clustering and UpSetR plots; Alyssa Shields, Avneet Brar and Andy Tran for technical assistance; Max Pottier and Gena Braun for instrumentation support. Research funding to the Castroverde Lab was provided by the NSERC Discovery Grant, Canada Foundation for Innovation, Ontario Research Fund, Faculty of Science start-up funds and Wilfrid Laurier University Internal Equipment. Vanessa Shivnauth was partially supported by a MITACS Research Training Award.

## Authors’ contributions

C.D.M.C conceptualized and supervised the study. V.S. identified the tomato *SlCBP60* genes and performed most of the experiments. S.P helped with the phylogenetic analyses, conducted the systemic gene expression analyses and performed the TM-score and PlantPAN analyses. E.M. completed the experimental replicates for the gene expression analyses. K.A. performed the structural predictions of the tomato SlCBP60 proteins. Everyone analyzed the data. V.S., S.P. and C.D.M.C. wrote the paper with input from all authors.

## Data availability

All data supporting the findings of this research are available within the main figures and supplementary materials.

## Supplementary Information

Supplementary Table 1. Table of qPCR primers

Supplementary Table 2. Table of SlCBP60 protein sequences

Supplementary Table 3. Pairwise TM-Score scores for the tomato and *Arabidopsis* CBP60 proteins

Supplementary Table 4. PlantPAN-predicted transcription factors that could bind the *SlCBP60* promoter regions

Supplementary Data 1. AlphaFold structures of the SlCBP60 proteins in PDB format

