## Supplementary figures and images for "Structural diversity and stress regulation of the plant immunity-associated CALMODULIN-BINDING PROTEIN 60 (CBP60) family of transcription factors in *Solanum lycopersicum* (tomato)"

### SOLYC01G100240_9B3D2_UNRELAXED_MODEL_5_RANK_1.PDB.png

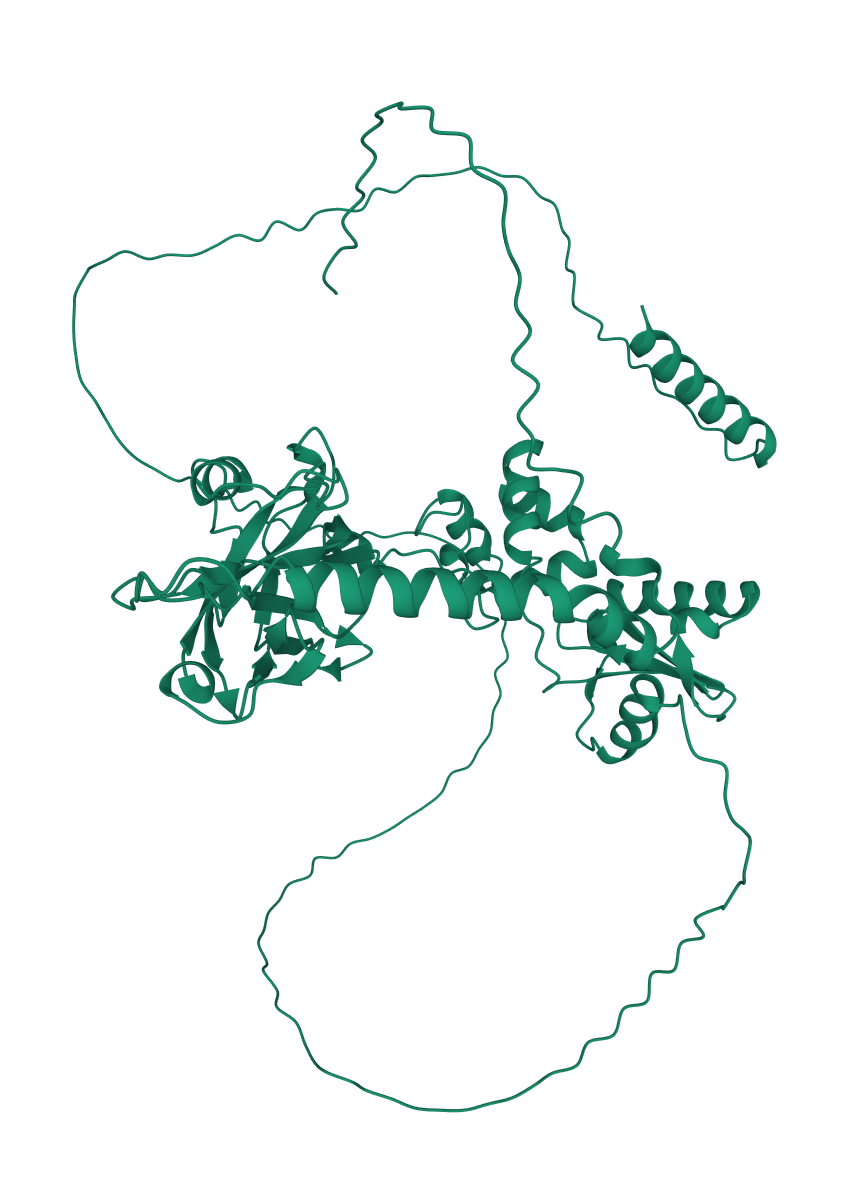

### SOLYC02G079040_1FB04_UNRELAXED_MODEL_3_RANK_1.PDB.png

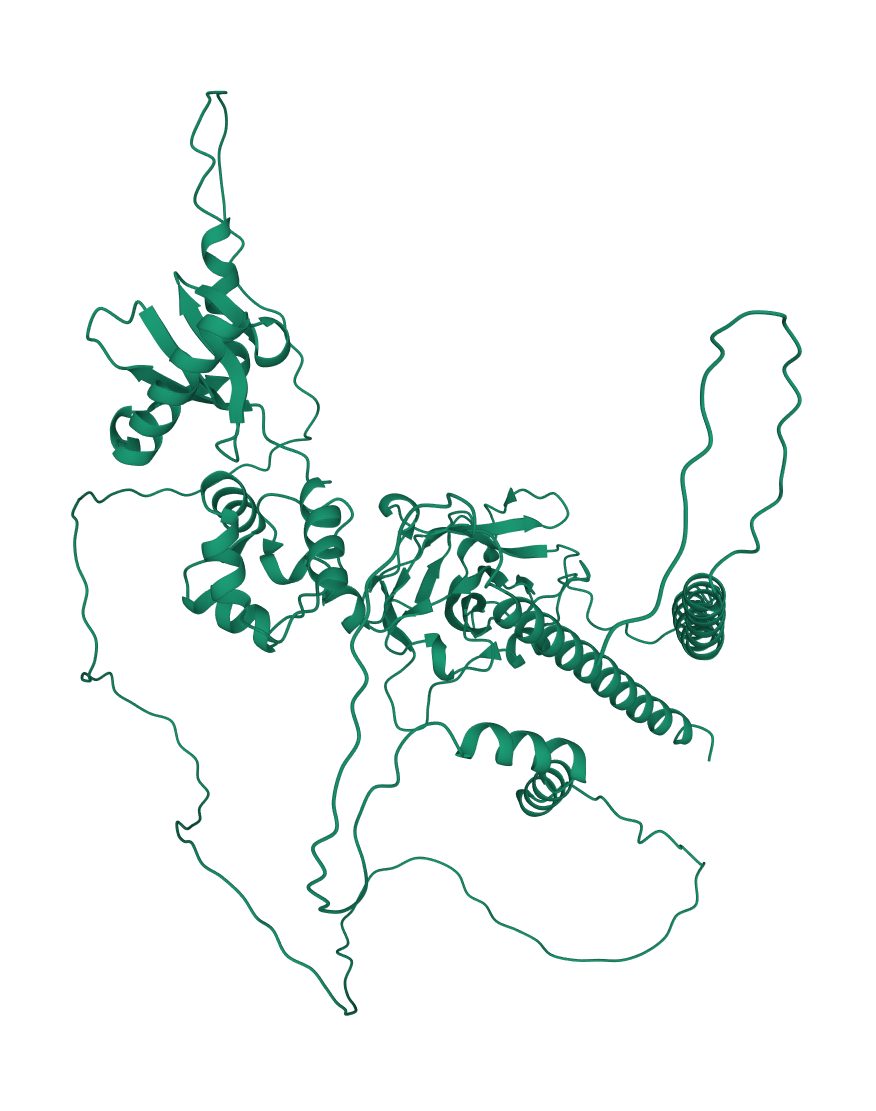

### SOLYC03G113920_FE1A3_UNRELAXED_MODEL_3_RANK_1.PDB.png

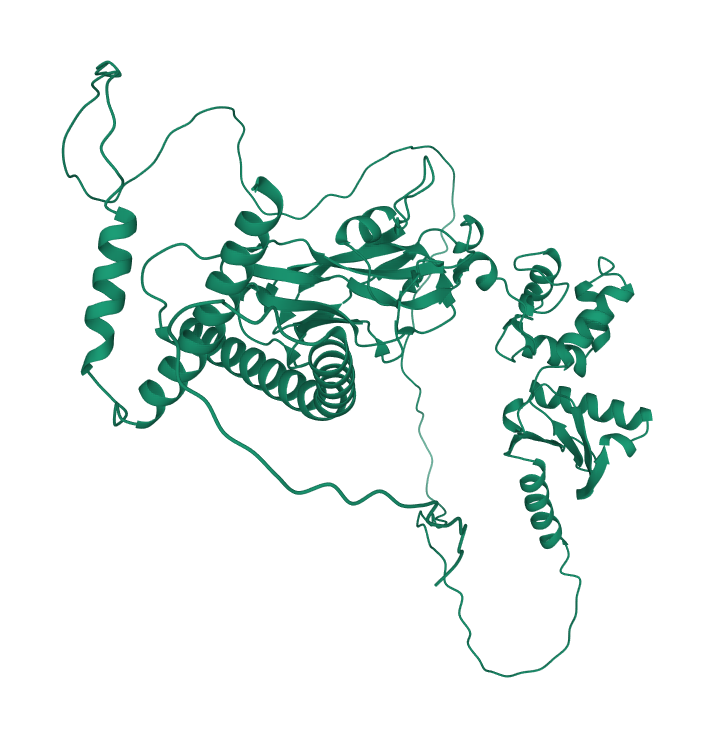

### SOLYC03G113940_BD418_UNRELAXED_MODEL_2_RANK_1.PDB.png

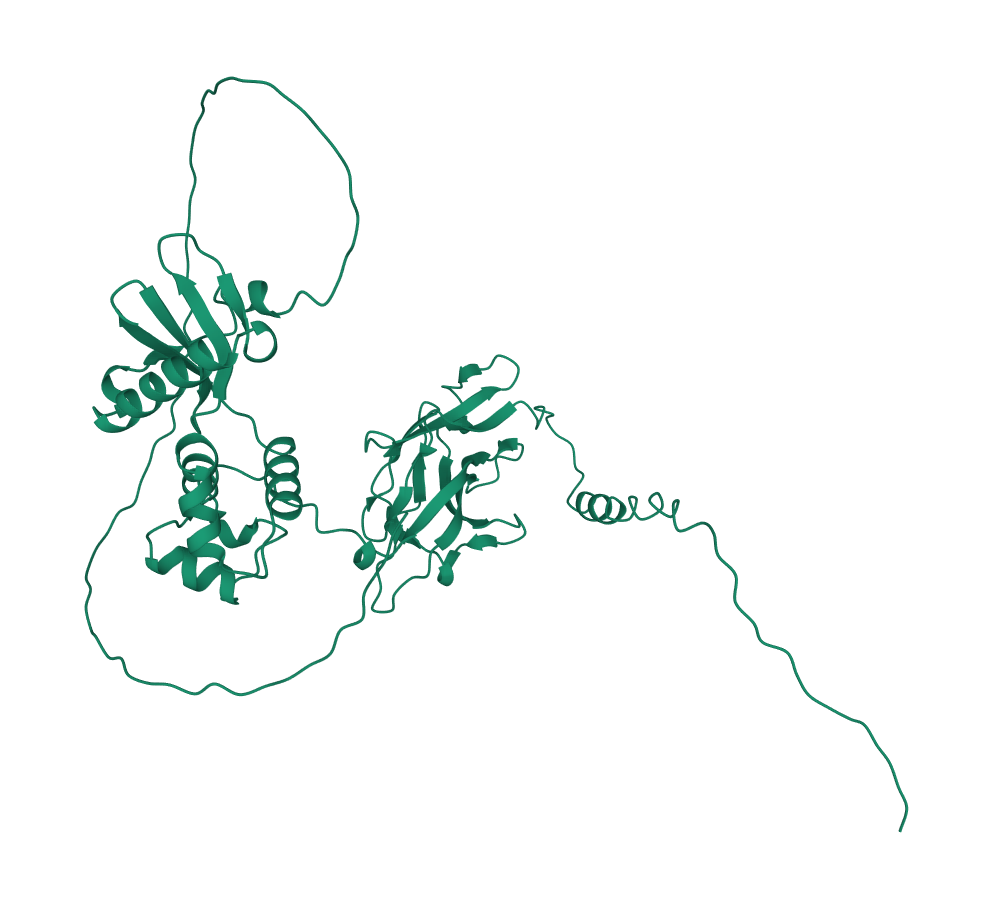

### SOLYC03G113950_62A9F_UNRELAXED_MODEL_5_RANK_1.PDB.png

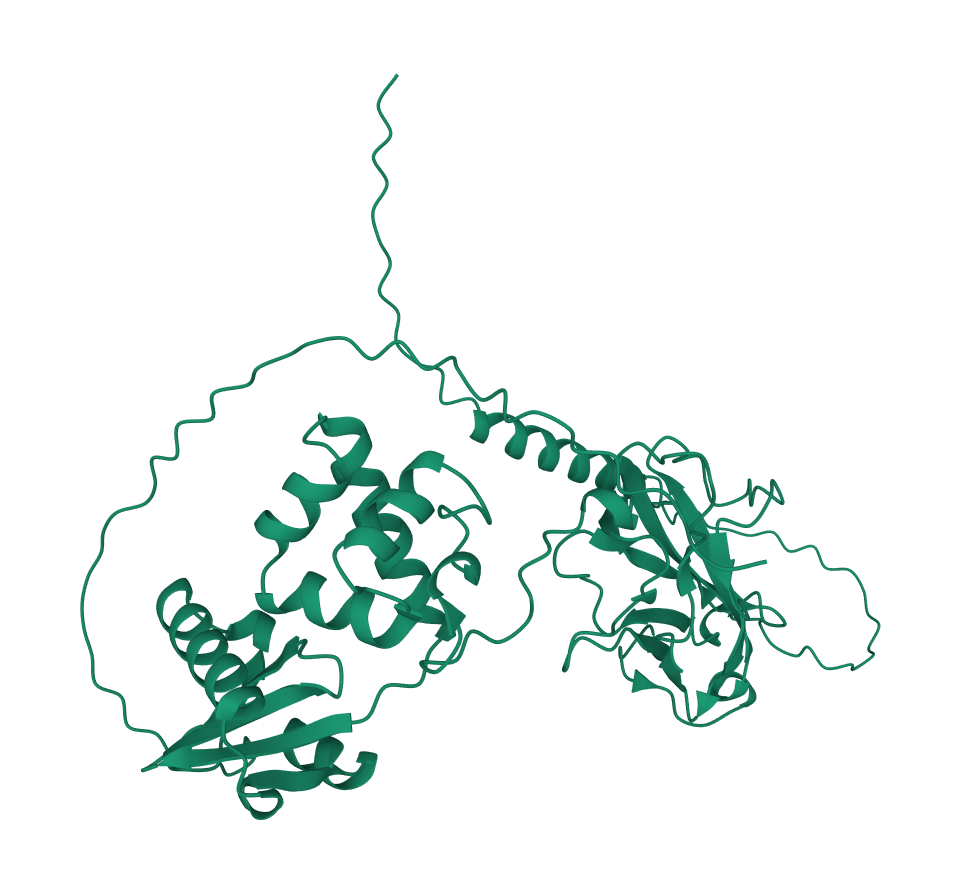

### SOLYC03G113960_BC04C_UNRELAXED_MODEL_2_RANK_1.PDB.png

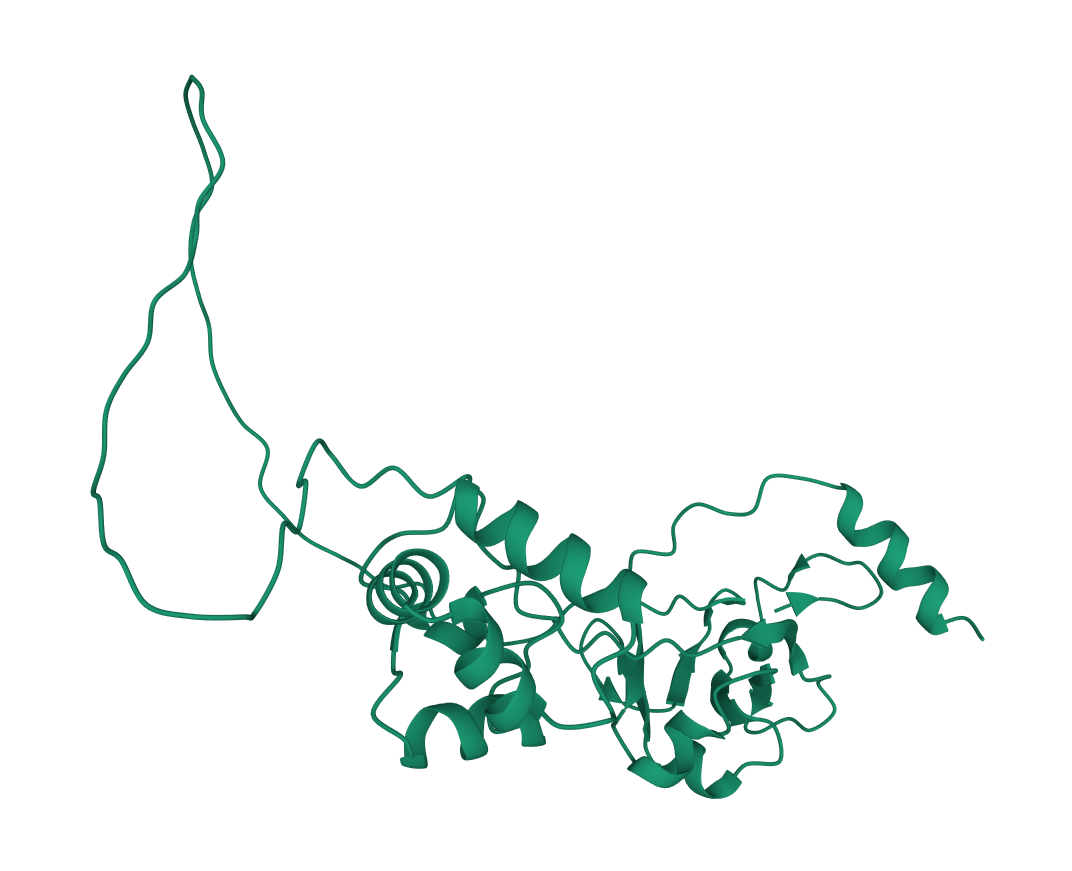

### SOLYC03G113970_28CF4_UNRELAXED_MODEL_5_RANK_1.PDB.png

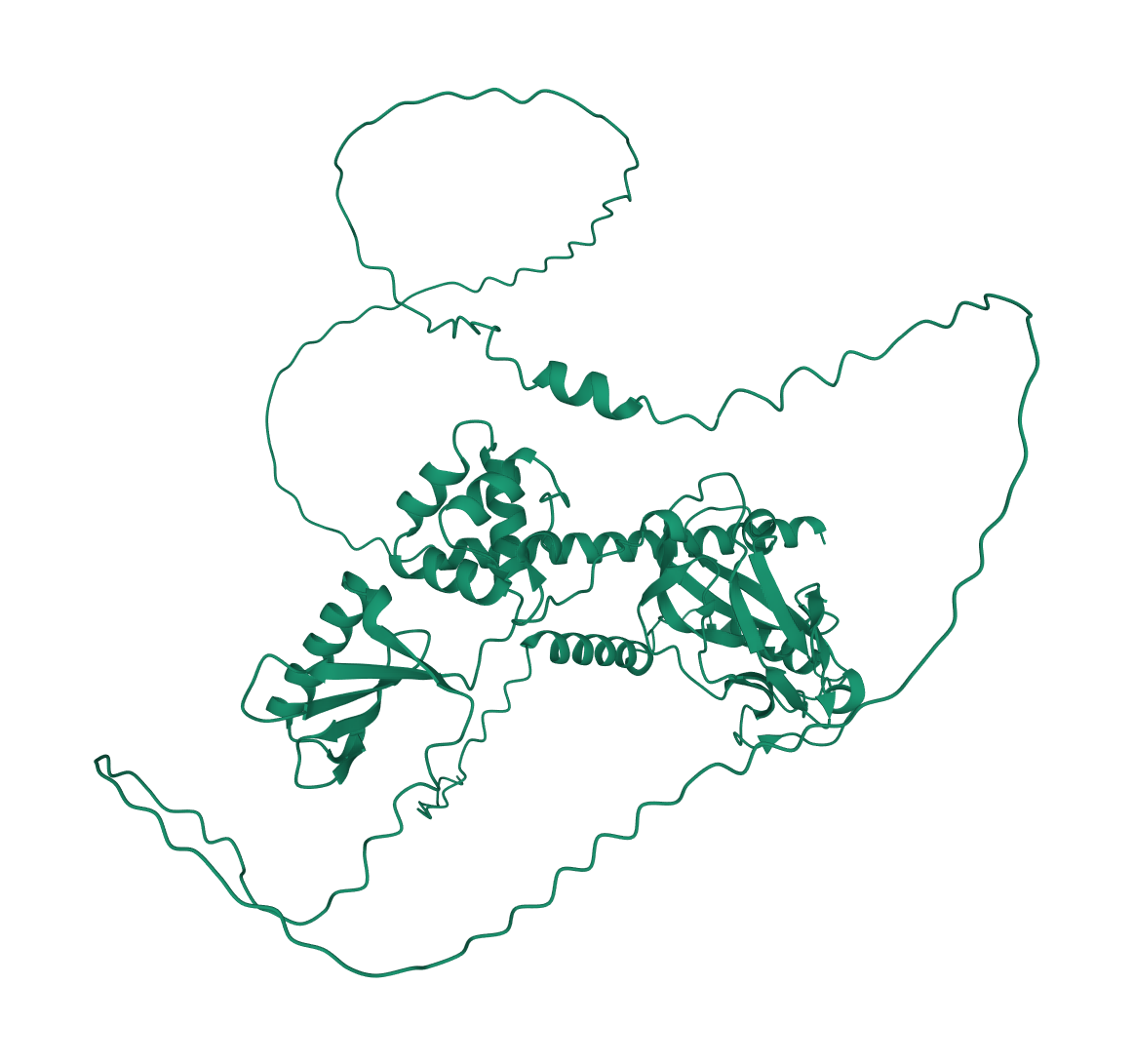

### SOLYC03G119250_F8A14_UNRELAXED_MODEL_5_RANK_1.PDB.png

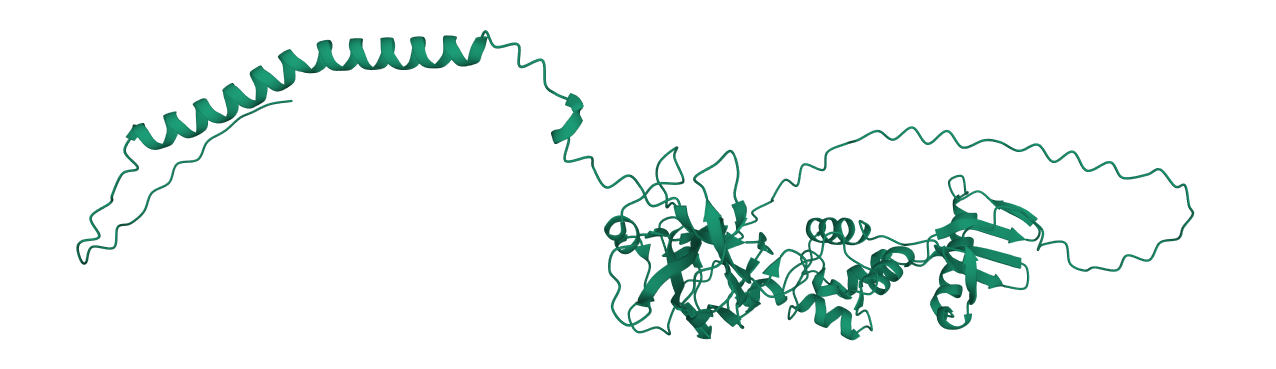

### SOLYC07G006830_0DCFF_UNRELAXED_MODEL_3_RANK_1.PDB.png

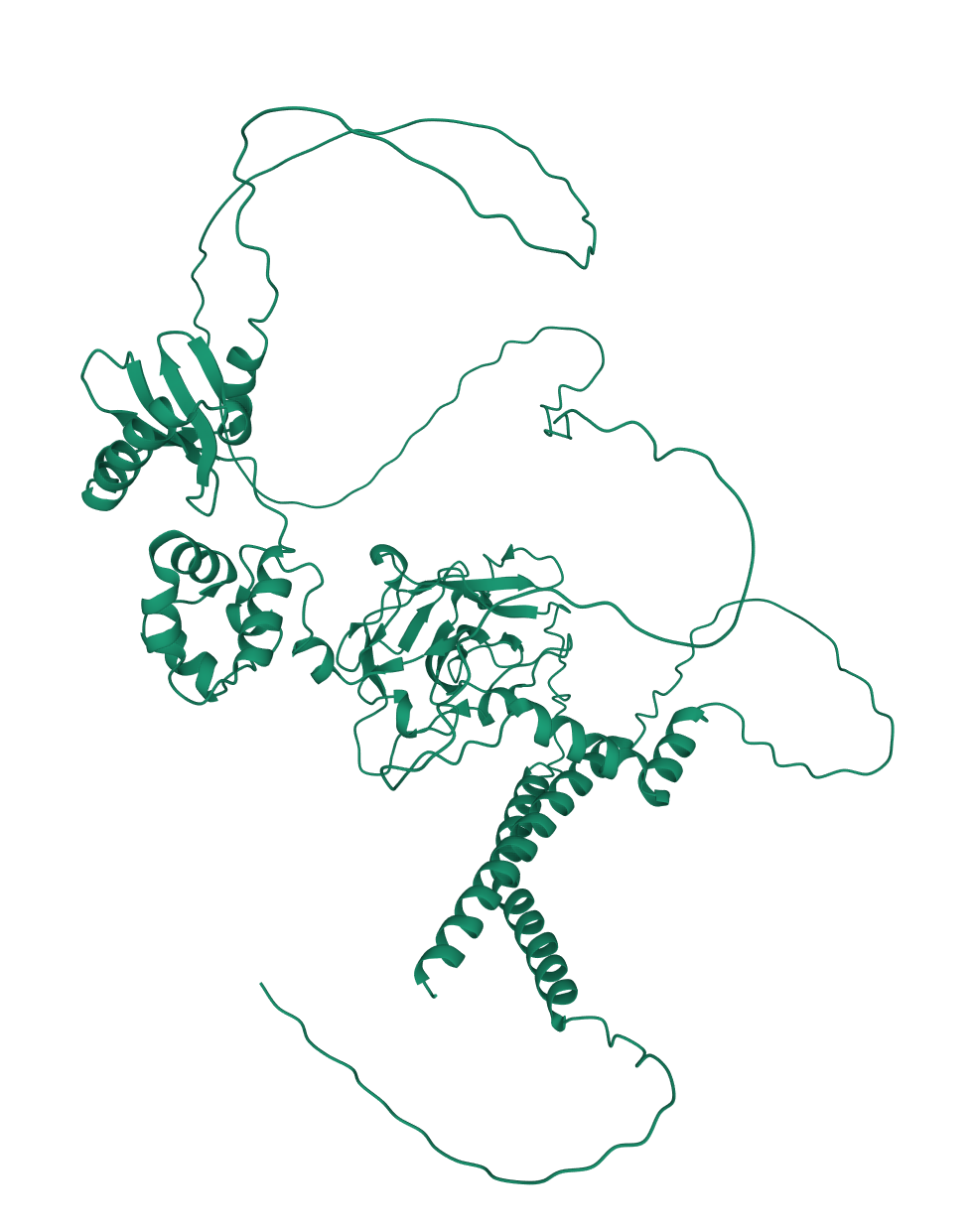

### SOLYC10G009210_A98DB_UNRELAXED_MODEL_3_RANK_1.PDB.png

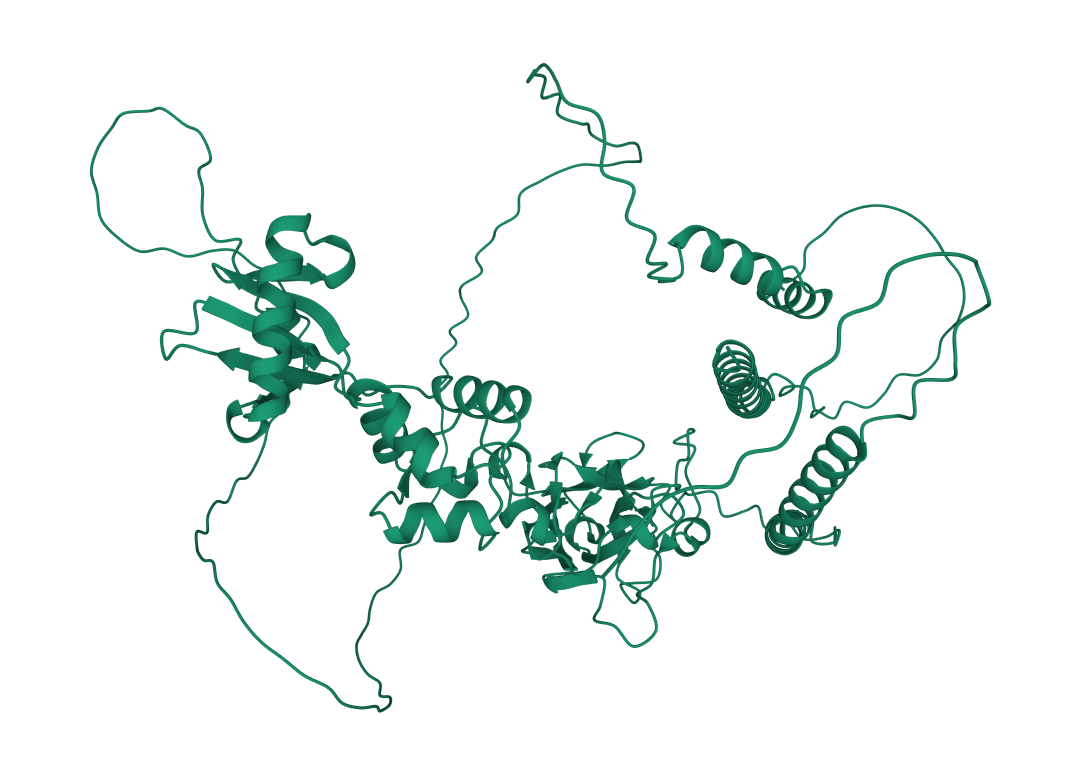

### SOLYC12G036390_99CA6_UNRELAXED_MODEL_5_RANK_1.PDB (1).png

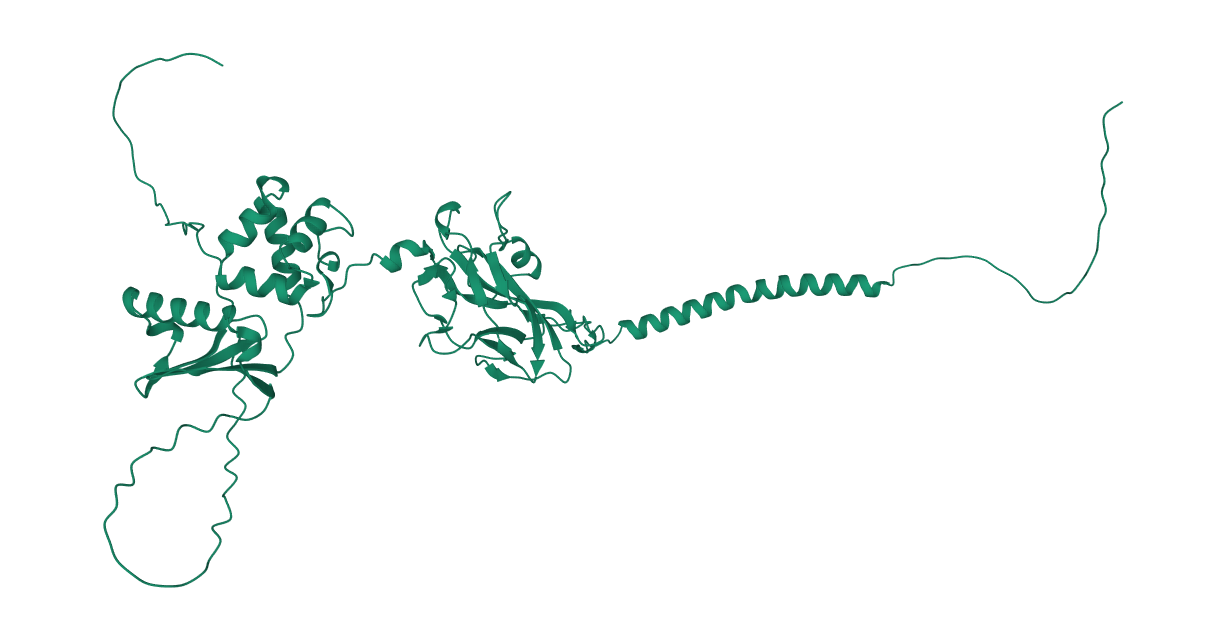
